# Neurogenin-2 Reprograms Human Microglial Lineage Cells into Neurons In Vitro and in Chimeric Brains

**DOI:** 10.64898/2026.03.19.712907

**Authors:** Mengmeng Jin, Ziyuan Ma, Rui Dang, Haiwei Zhang, Haipeng Xue, Steven Finkbeiner, Ying Liu, Peng Jiang

## Abstract

Progressive neuronal loss is a hallmark of many neurological disorders, yet the adult human brain has a limited capacity for endogenous neuronal replacement. Direct neuronal reprogramming represents an alternative strategy for generating new neurons. Microglia, the brain’s resident immune cells, are uniquely positioned as candidate cellular substrates due to their abundance, self-renewal capacity, high motility, and rapid recruitment to sites of injury. Here, using live-cell imaging and electrophysiological recordings, we show that human pluripotent stem cell (hPSC)–derived primitive macrophage progenitors (PMPs) and their microglial derivatives exhibit neuronal reprogramming competence. Inducible expression of NEUROG2 in hPSC-derived PMPs drives acquisition of neuronal morphology, sequential expression of early and mature neuronal markers, organization of synaptic proteins, and functional excitability characterized by action potential firing. Single-nucleus RNA sequencing reveals a continuous, directionally ordered reprogramming trajectory marked by suppression of myeloid transcriptional programs, progression through intermediate remodeling states, and progressive activation of neuronal gene regulatory networks, consistent with a regulated lineage conversion rather than partial identity switching. Using a xenotransplantation-based human microglia chimeric brain model, we further demonstrate that inducible NEUROG2 expression reprograms donor-derived human microglia toward a neuronal identity *in vivo*. Together, these findings establish human microglial lineage cells as a previously unexplored substrate for neuronal reprogramming, providing a conceptual framework for microglia-based strategies aimed at neuronal replacement and neural repair.

## Introduction

Progressive and irreversible neuronal loss is a defining feature of many central nervous system (CNS) disorders, including neurodegenerative diseases, stroke, and traumatic brain injury ^1–4^. Because the adult mammalian CNS exhibits limited regenerative capacity, strategies capable of replacing lost neurons and restoring functional neural circuits remain a long-standing therapeutic goal ^5,6^. Although stimulation of endogenous neural stem cells (NSCs), neuroprotective interventions, and transplantation of human stem cell-derived neurons have shown promise, each faces persistent challenges related to integration, survival, spatial distribution, and long-term functional efficacy ^7–9^. A conceptually attractive alternative that has gained attention is the in-situ reprogramming of resident glial cells into functional neurons through the delivery of neurogenic transcription factors ^10^. The promise of direct neuronal reprogramming of macroglia, including astroglia and oligodendroglia, has been well-established across various *in vivo* disease models, demonstrating impressive results in neuronal restoration, functional integration, and subsequent behavioral recovery ^11–22^.

Microglia represent an especially intriguing yet challenging cellular substrate for neuronal reprogramming. They are abundant, self-renewing, highly motile, and capable of brain-wide migration with rapid recruitment to sites of CNS injury ^23,24^. These distinctive properties position microglia as a biologically compelling and strategically advantageous population for in situ neuronal replacement. Nevertheless, microglia-to-neuron (MtN) reprogramming has become controversial, with reports describing discrepant outcomes, ranging from successful neuronal conversion to microglial apoptosis, following ectopic expression of the transcription factor NeuroD1 ^25–27^. Importantly, foundational work on cross-lineage neural fate conversion has firmly established both the feasibility and mechanistic basis of transcription factor–driven cell identity switching ^10,28,29^, suggesting that careful selection of transcriptional drivers and contextual conditions will be critical for efficient neuronal generation from microglia. Moreover, nearly all prior MtN attempts have been conducted using rodent microglia or macrophages ^25,30,31^. Given substantial species-specific differences between mouse and human microglia, encompassing transcriptional identity and signaling pathways ^32–34^, whether human microglial lineage cells can be directly converted into neurons remains a critical unresolved question. In contrast to transplanted human neural progenitor cells or their neuronal derivatives, which typically remain confined near injection sites, studies from our group and others have shown that human pluripotent stem cell (hPSC)–derived microglia not only exhibit a unique capacity for extensive migration following transplantation, but also actively respond to and are rapidly recruited to sites of brain injury ^35–38^. Successful reprogramming of hPSC-derived microglia into neurons could therefore establish a distinct therapeutic paradigm, in which xenografted microglia equipped with inducible neuronal reprogramming programs function as programmable neuron-producing units to enable neuronal replacement and neural repair.

Transcription factor–driven reprogramming is particularly powerful, as proneural factors associated with neuroectodermal lineage specification can robustly activate neuronal gene regulatory programs. Among them, neurogenin 2 (NEUROG2 or NGN2) is a basic helix–loop–helix (bHLH) transcription factor that plays a pivotal role during embryonic neurogenesis by specifying neuronal fate and driving differentiation. It promotes rapid cell-cycle exit, suppresses glial identity, binds to E-box motifs within neuronal gene enhancers, directly activates downstream neuronal transcription factors such as NEUROD1 and BRN2, and contributes to neuronal migration and subtype specification ^39,40^. NEUROG2 has been shown to robustly reprogram fibroblasts, astrocytes, oligodendrocyte progenitor cells, and even certain tumor cells into functional neurons ^41–45^. In this study, we leveraged CRISPR-engineered and doxycycline-inducible NEUROG2 hPSC lines ^46^, high-purity derivation of yolk-sac primitive macrophage progenitors (PMPs) from hPSCs ^47,48^, and *in vivo* lineage tracing using a human microglia chimeric brain model ^38,49^ to interrogate NEUROG2-driven MtN conversion. Using a rigorously controlled human *in vitro* system, we tested the hypothesis that NEUROG2 triggers a coordinated transcriptional and functional transition from myeloid to neuronal identity. By integrating electrophysiological measurements with single-cell transcriptomic analyses, we resolved the cellular trajectories underlying reprogramming and defined molecular states associated with successful lineage conversion. Furthermore, using an *in vivo* human microglia chimeric brain model ^38,49^, we provide proof-of-concept evidence that xenografted hPSC-derived microglia can be converted into neurons in the brain. Collectively, these findings establish a robust human experimental framework that demonstrates MtN fate plasticity and provide foundational support for a fundamentally new strategy in human neuronal replacement and regenerative repair.

## Results

### NEUROG2 reprograms human microglial lineage cells into neurons with hallmark morphological and functional properties

To determine whether human microglial lineage cells can undergo neuronal lineage conversion, we utilized CRISPR–Cas9–mediated gene editing to establish a doxycycline (Dox)–inducible NEUROG2 expression system in two hESC lines, H1 and H9 (Fig. S1A-B). Subsequently, we differentiated NEUROG2 hESCs into primitive macrophage progenitor (PMP) cells. A schematic overview of the experimental paradigm is shown in Fig. 1A, illustrating the induction of NEUROG2 expression for 10 days, followed by sequential morphological, molecular, electrophysiological, and transcriptomic analyses at defined timepoints. A critical requirement for interpreting lineage conversion is to verify that the initial cellular population does not contain pre-existing neural lineage contaminants. To address this, we first characterized the PMP-derived population prior to reprogramming. As shown in Fig. S1C, owing to the unique PMP differentiation protocol of supernatant harvest of PMPs and consistent with previous reports ^50,51^, the resulting cell population is pure, with PMPs robustly expressing the hematopoietic lineage markers CD43 and CD235, thereby confirming their myeloid origin. In contrast, markers of neural stem cells (SOX2), astrocytes (SOX9, GFAP), and oligodendrocyte lineage cells (OLIG2) were undetectable (Fig. S1C–D). Consistent with the immunostaining results, quantitative PCR revealed undetectable expression of SOX2, DCX, OLIG2, and GFAP, whereas GAPDH was robustly expressed (Fig. S1D). These findings verify that the starting population lacks neural lineage identity and therefore exclude the possibility that the neurons observed following NEUROG2 induction arise from contaminating neural progenitors. Then, we collected these PMPs and transferred them into the neuron differentiation (ND) medium containing Dox and Rho-associated protein kinase (ROCK) inhibitor Y-27632, which promotes cell survival during neuronal lineage conversion ^52^ (Fig. 1A). We next asked whether ectopic induction of NEUROG2 could redirect PMPs toward activation of a neuronal fate. We first observed that by day 5, cells began to elongate and extend neurite-like processes. By day 10, the cultures displayed an increasingly complex arborization pattern, and by day 28, a dense neuronal network with extensive process outgrowth was evident. These changes were not observed in non-induced controls, indicating that neuronal-like morphological acquisition is NEUROG2 dependent (Fig. 1B). At day 10 post-induction, DOX-treated cultures remained negative for glial lineage markers, including SOX9, OLIG2, and GFAP, while control cultures also showed no evidence of neural marker expression (Fig. S1E). Instead, neuronal marker expression selectively emerged in Dox-treated cultures, and hematopoietic marker expression (CD235) diminished in reprogramming cells (Fig. S1E), consistent with a fate transition. Together, these observations indicate that NEUROG2 reprogramming does not result from spontaneous neural differentiation, nor does it broadly activate non-neuronal neural programs. Rather, the data support a directed neuronal lineage conversion from human microglial lineage cells.

**Fig 1.**
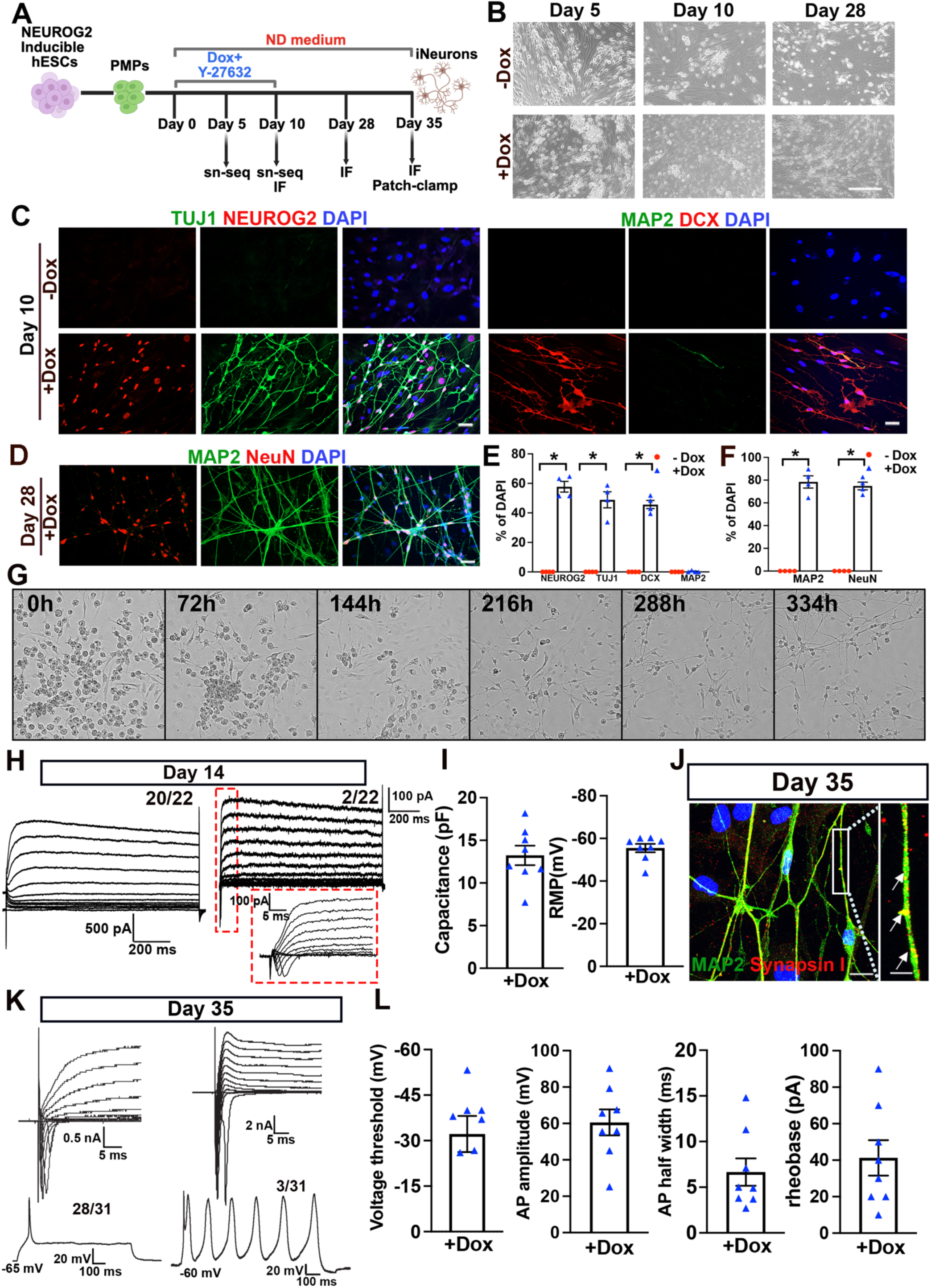
NEUROG2 induces neuronal reprogramming and functional maturation of PMPs *in vitro*. A. Schematic of the experimental design. B. Phase-contrast images showing morphological progression of no Dox and Dox-treated cultures at Days 5, 10, and 28. Scale bars: 750 μm. C. Representative images of NGN2, TUJ1, DCX, and MAP2. Scale bars: 20 μm. D. Representative images of MAP2 and NeuN. Scale bars: 20 μm. E–F. Quantification of neuronal marker expression at Day 10 (E) and Day 28 (F). (n=4, each experiment was repeated three times). Data are presented as mean ± SEM; each point represents an independent biological replicate. Data were analyzed by a two-tailed unpaired t-test. Statistical significance determined as indicated; \**P < 0.05*. G. Time-lapse phase-contrast imaging illustrating the dynamic morphological progression during reprogramming. Time is shown in hours post-induction. H. Representative voltage-clamp recordings from +Dox cells at day 14 showing voltage-dependent inward and outward currents. Zoom in (red dashed rectangle): fast activation/inactivation voltage-dependent sodium currents. I. Quantification of passive membrane properties at day 14, including membrane capacitance and resting membrane potential (RMP). J. Representative images of MAP2-positive dendritic arbors with punctate Synapsin I localization. Scale bars: 20 μm and 5 μm in the original and enlarged images, respectively. K. Representative voltage-clamp and current-clamp recordings from Day 35 +Dox cells. L. Quantification of intrinsic excitability parameters at day 35, including voltage threshold, rheobase, action potential amplitude, and action potential half-width. Data are presented as mean ± SEM; each point represents an individual recorded cell.

Next, we examined whether these morphological changes were accompanied by molecular signatures of neuronal lineage acquisition. At day 10 following induction, immunofluorescence analysis demonstrated robust expression of early neuronal markers, including the pan-neuronal cytoskeletal marker TUJ1 and the neuronal migration/immaturity marker doublecortin (DCX), exclusively in Dox-treated cultures (Fig. 1C). NEUROG2 protein was readily detectable in Dox-treated cells, confirming efficient induction of the transgene. In contrast, control cultures lacked detectable expression of neuronal lineage markers. Quantification confirmed that a substantial proportion of total cells expressed NEUROG2, TUJ1, and DCX at this reprogramming stage (Fig. 1E), supporting the establishment of early neuronal identity. To examine whether these reprogrammed cells further progressed toward mature neuronal states, we assessed expression of late neuronal markers at later timepoints. By day 28, Dox-treated cultures exhibited widespread expression of MAP2, a dendritic neuronal marker, and NeuN, a nuclear marker of neuronal identity, accompanied by highly elaborated neuritic networks (Fig. 1D). Control cultures remained negative for neuronal markers. Quantitative analysis demonstrated that a large fraction of DAPI-positive cells expressed MAP2 and NeuN at this stage (Fig. 1F), indicating continued maturation of neuronal features over time. To directly visualize the dynamics of cellular remodeling, longitudinal time-lapse imaging was performed throughout the reprogramming process. PMP cultures initially exhibited small, round, loosely adherent cells at the onset of induction. Over the course of several days, NEUROG2-induced cells progressively increased substrate adherence, elongated, and extended polarized neurite-like processes. By approximately two weeks post-induction, cells formed interconnected networks with a pronounced neuronal-like morphology, a change not observed in the no Dox-treated cultures groups (Fig. 1G, and Video S1 and S2). Collectively, these data demonstrate that NEUROG2 expression initiates a progressive neuronal reprogramming trajectory, driving the acquisition of neuronal identity in human microglial lineage cells. This process is characterized by stepwise morphological transformation and the sequential acquisition of early and mature neuronal marker expression.

To determine whether NEUROG2-induced human microglial lineage–derived induced neurons (iNs) not only express neuronal identity markers but also acquire structural and functional properties consistent with neuronal maturation, we performed whole-cell patch-clamp electrophysiology at different stages following reprogramming. Prior to reprogramming, PMPs exhibited minimal voltage-dependent currents under voltage-clamp conditions, lacking fast inward or delayed outward currents characteristic of voltage-gated sodium and potassium channels (Fig. S1F). Quantification of passive membrane properties revealed that PMPs displayed relatively large membrane capacitance and depolarized resting membrane potentials, consistent with a macrophage-like, non-excitable cellular state (Fig. S1G). These baseline measurements confirm that PMPs do not possess latent neuronal electrophysiological properties prior to the induction of NEUROG2. At day 14 after Dox treatment, the majority of NEUROG2-induced cells (20 of 22 recorded cells) exhibited prominent voltage-dependent outward currents under voltage-clamp conditions, while a smaller fraction (2 of 22) displayed fast inward currents consistent with the emergence of sodium conductance (Fig. 1H). These findings indicate that ion channel expression is initiated early during reprogramming but continues to mature over time. Passive membrane property analysis revealed that induced cells exhibited membrane capacitance and resting membrane potentials (RMP) within the ranges expected for developing neurons at day 14 (Fig. 1I). By day 35, NEUROG2-induced iNs exhibited extensive MAP2-positive dendritic arbors with punctate Synapsin I localization along neurites (Fig. 1J), supporting the formation of synaptic protein assemblies at later stages of maturation. Electrophysiological recordings at day 35 demonstrated further maturation of intrinsic excitability. Under voltage-clamp conditions, iNs displayed robust inward and outward voltage-gated currents (Fig. 1K). In current-clamp mode, the majority of recorded cells (28 of 31) generated action potentials in response to depolarizing current injection, with a subset (3 of 31) capable of firing repetitive action potentials (Fig. 1K). Quantitative analysis revealed well-defined voltage thresholds, substantial action potential amplitudes, measurable half-widths, and rheobase currents within the expected range for developing human neurons (Fig. 1L). Collectively, these data demonstrate that NEUROG2-induced human microglial lineage–derived iNs undergo a progressive maturation process characterized by the sequential acquisition of ion channel activity, intrinsic excitability, and synaptic protein organization. The delayed emergence of repetitive firing and synaptic marker clustering suggests ongoing functional refinement over extended culture periods, consistent with a gradual transition toward a more mature neuronal state.

### Single-nucleus RNA sequencing identifies progressive transcriptional states linking microglial lineage identity to neuronal fate

To define the molecular dynamics underlying human microglial lineage cells-to-neuron reprogramming, we performed single-nucleus RNA sequencing (snRNA-seq) on PMPs cultured in the ND medium for 5 or 12 days, with or without Dox induction. NEUROG2 transcripts were detected exclusively in Dox-treated cultures, confirming inducible and condition-specific expression (Fig. 2A). Leiden clustering of UMAP-projected nuclei identified seven transcriptionally distinct clusters (Fig. 2B), comprising two PMP clusters, one transitioning PMP cluster, two intermediate clusters, one neuronal cluster (Fig. 2C), and one small cluster containing only 47 cells (0.05% of total cells) that showed low similarity to known cell types (Fig. S2A). This minor cluster likely reflects an artifact of culturing PMPs under neuronal differentiation conditions and was therefore excluded from subsequent analyses. Cell-type proportion analysis revealed a marked enrichment of neurons in the 12-day Dox-treated cultures, accompanied by a corresponding reduction in PMP populations (Fig. 2D). We also detected a small neuronal cluster comprising 5.8% of cells in the 12-day no-Dox condition. However, these cells did not express canonical neuronal marker genes (Fig. 2E). Differentially expressed gene (DEG) analysis comparing cells assigned to the neuronal cluster between Dox-treated and untreated cultures revealed that genes upregulated in the Dox-treated group were significantly enriched for gene ontology (GO) terms related to neuronal development, synaptic function, and membrane potential regulation. In contrast, no neuron-related GO terms were enriched among genes upregulated in the no-Dox group (Fig. 2F). We speculate that human PSC–derived PMPs retain higher cellular plasticity than fully mature cells and are therefore more responsive to in vitro neuronal culture conditions. This heightened responsiveness may result in partial activation of neuron-associated transcriptional programs, which in turn could lead to misclassification of these cells within neuronal clusters during transcriptomic analyses. To examine this possibility, we further performed whole-transcriptome similarity analysis comparing neuronal-cluster cells with other cell populations. Cells from the no-Dox cultures exhibited low transcriptomic similarity to the overall bona fide neuronal population, with similarity indices instead closer to those of intermediate reprogramming states (Fig. 2G). Consistent with this interpretation, immunostaining failed to detect any cells positive for neuronal protein markers in no-Dox cultures *in vitro* (Fig. 1C), providing independent validation that these cells reflect transcriptional responses of PMPs to neuronal culture conditions rather than spontaneous transdifferentiation. Notably, to our knowledge, spontaneous cross-lineage transdifferentiation from the myeloid to the neural lineage has not been reported to occur solely as a consequence of altered culture conditions and is therefore considered unlikely. Consistent with prior reports of NEUROG2 activity, reprogrammed neurons expressed markers of excitatory, but not inhibitory, neuronal identity (Fig. 2H).

**Fig 2.**
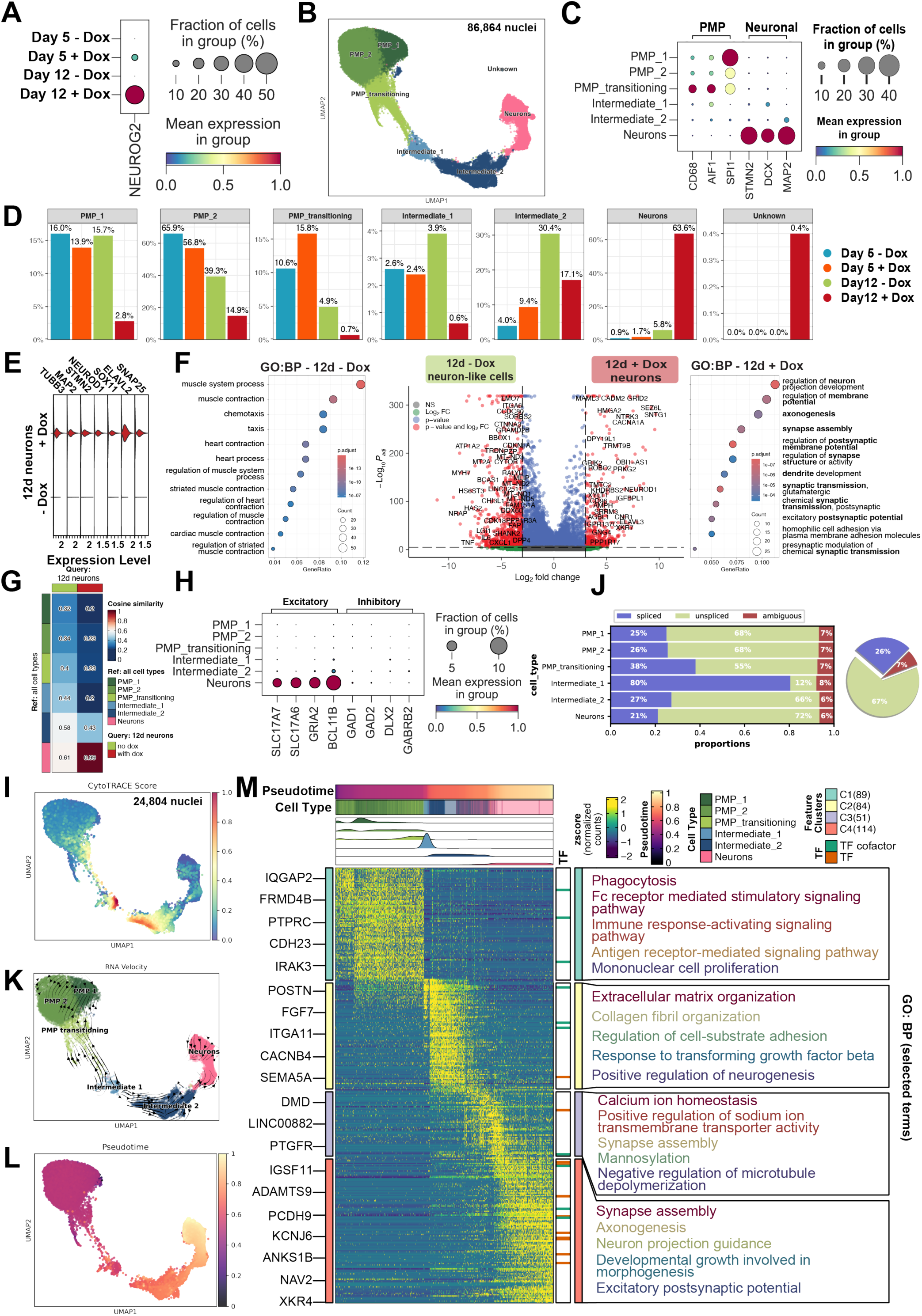
Single-nucleus RNA sequencing of human microglial lineage cells during NEUROG2-mediated reprogramming. A. Dot plot showing NEUROG2 expression across 4 experimental groups. B. UMAP visualization of 86,864 single nuclei collected across 4 experimental groups. C. Dot plot showing expression of lineage markers across major Leiden clusters. D. Bar plot showing the proportion of cells from each experimental group across annotated cell types. E. Stacked violin plot showing expression of canonical neuronal marker genes within the neuronal cluster at 12-day +Dox and -Dox conditions. F. Volcano plot and dot plots showing differentially expressed genes (DEGs) and associated GO enrichment between +Dox and -Dox groups at the day 12 time point. G. Heatmap showing cosine similarity indices comparing day 12 +Dox and −Dox neuronal populations with other identified cell types. H. Dot plot showing expression of canonical excitatory and inhibitory neuronal lineage markers across identified clusters. I. UMAP visualization of differentiation potential inferred from 24,804 nuclei from day 12 samples. J. Bar plot and pie chart showing the proportions of spliced and unspliced RNA across individual cell types and the overall population, respectively. K. UMAP colored by cell type, overlaid with unified RNA velocity streamlines and individual cell velocity vectors. L. UMAP visualization showing velocity-inferred pseudotime progression. M. Dynamic heatmap (left) showing DEG expression changes along pseudotime and corresponding GO term enrichment analysis (right).

To characterize reprogramming dynamics, we inferred developmental potential across cell states. Differentiation potential increased in the transitioning PMP cluster, peaked in the intermediate clusters, and declined in the neuronal cluster (Fig. 2I), suggesting that reprogramming passes through a highly plastic, progenitor-like phase. RNA velocity analysis further revealed state-dependent transcriptional kinetics, with a higher proportion of spliced relative to unspliced transcripts in transitioning and intermediate populations compared with initial PMP and terminal neuronal states, suggesting slowed or attenuated transcriptional dynamics (Fig. 2J). Projecting velocity vector fields onto the UMAP embedding revealed that velocity vectors in the “Transitioning PMP” and “Intermediate 1” clusters were incoherent or reversed toward the PMP state (Fig. 2K). This suggests a strong transcriptional barrier during the mesoderm-to-ectoderm lineage conversion. However, cells that traversed this barrier entered “Intermediate 2,” where velocity vectors flowed coherently toward the neuronal cluster. Pseudotime analysis confirmed a continuous trajectory from PMPs through intermediate stages to neurons (Figs. 2L, S2B). In order to further examine transcriptional programs associated with this trajectory, we performed k-means clustering of genes dynamically regulated along velocity pseudotime. Dynamic gene cluster 1 was enriched in PMP populations and associated with GO terms related to phagocytosis and immune activation, consistent with myeloid identity. Dynamic gene clusters 2 and 3 were enriched in intermediate stages and associated with GO terms including extracellular matrix organization and ion homeostasis, reflecting transcriptional programs linked to structural remodeling and signaling changes during lineage transition. Dynamic cluster 4 was selectively enriched in the neuronal cluster and associated with neuronal processes, including synapse assembly, axonogenesis, and neuronal morphogenesis (Fig. 2M). Together, these data demonstrate that NEUROG2 induces a transcriptionally continuous and directional reprogramming trajectory from a myeloid microglial lineage toward a neuron-like state.

### NEUROG2-mediated reprogramming is accompanied by stepwise rewiring of transcription factor regulatory networks

To dissect the gene regulatory networks (GRNs) driving this conversion, we performed *de novo* transcription factor enrichment and regulon inference (Figs. 3A, S2C). PMP clusters were characterized by high activity of canonical myeloid regulons, including SPI1, CEBPB, and STAT family members. The first intermediate cluster showed enrichment of regulons associated with stress and adaptive responses, including CREB3, FOXO4, JUND, and DDIT3. In the second intermediate cluster, regulons linked to chromatin organization and neuronal differentiation, such as GATA6, NFIB, and FOXP1, became prominent. In neuronal clusters, direct downstream targets of NEUROG2, most notably HES6 ^53,54^, emerged among the top regulons, together with additional neurogenic regulators including SOX11 and NFIA. To systematically characterize combinatorial regulation, we assessed pairwise regulon similarity using the connection specificity index (CSI). This analysis organized 223 regulons into six major modules (Fig. 3B). Module 3 was related to macrophage and PMP identity regulators, including ATF3, SPI1, and IRF family members, whereas Module 2 contained direct targets of NEUROG2 (HES6, EBF2) ^53^, neurogenic regulons (SOX4, SOX11), and patterning genes (HOXA2/3, LHX2). Module 4 exhibited transient enrichment in transitioning and intermediate populations, with reduced activity in neuronal clusters (Fig. S2J). This module represents a reprogramming barrier characterized by transcriptional conflict between myeloid and neuronal programs. It is enriched for stress-adaptive factors (ATF4, XBP1), neuronal repressors (REST, HES1), chromatin remodelers associated with plasticity (POU5F1, KLF family), and also pro-neuronal and transcriptional organizing factors (FOXP1), suggesting that neuronal differentiation is temporarily restrained to allow for chromatin stabilization at the intermediate stages. RNA velocity analysis supports this model: cells enriched for Module 4 displayed bidirectional velocity vectors, with some reverting toward the PMP state and others progressing toward neuronal commitment. Once cells overcame the reprogramming barrier, we observed coordinated repression of the neuronal repressor HES1, accompanied by the concomitant upregulation of HES6 and NFIA, both of which have been previously shown to function as negative regulators of HES1 activity ^55^ (Fig. 3C). This pattern suggests a synergistic regulatory mechanism that facilitates the release of HES1-mediated neuronal fate repression. Consistent with this transition, neuronal gene expression programs became progressively activated, reflecting enhanced neuronal lineage commitment. In parallel, increased activity of the transcriptional regulator YY1 (Fig. 3C), a known facilitator of cell fate reprogramming ^56^, further reinforced neuronal transcriptional programs, indicating a role in stabilizing the neuronal identity following successful reprogramming. Collectively, these data identify Module 4 as a critical checkpoint where cells must resolve transcriptional stress and repress premature differentiation to successfully initiate the neuronal program. Using the CSI, we constructed a regulon association network to resolve higher-order regulatory relationships during reprogramming (Fig. S2D). Hub regulons were identified based on degree centrality, revealing a subset of highly connected regulators that occupied central positions within the network (Figs. 3E-F). These hub regulons exhibited strong and specific co-association patterns, suggesting coordinated regulatory roles in controlling transcriptional state transitions during neuronal reprogramming. Together, these analyses demonstrate that NEUROG2-induced microglial reprogramming is accompanied by extensive remodeling of transcriptional regulatory networks. Rather than a binary switch, reprogramming proceeds through intermediate regulatory states marked by coordinated suppression of myeloid programs and progressive activation of neuronal transcriptional circuits.

**Fig 3.**
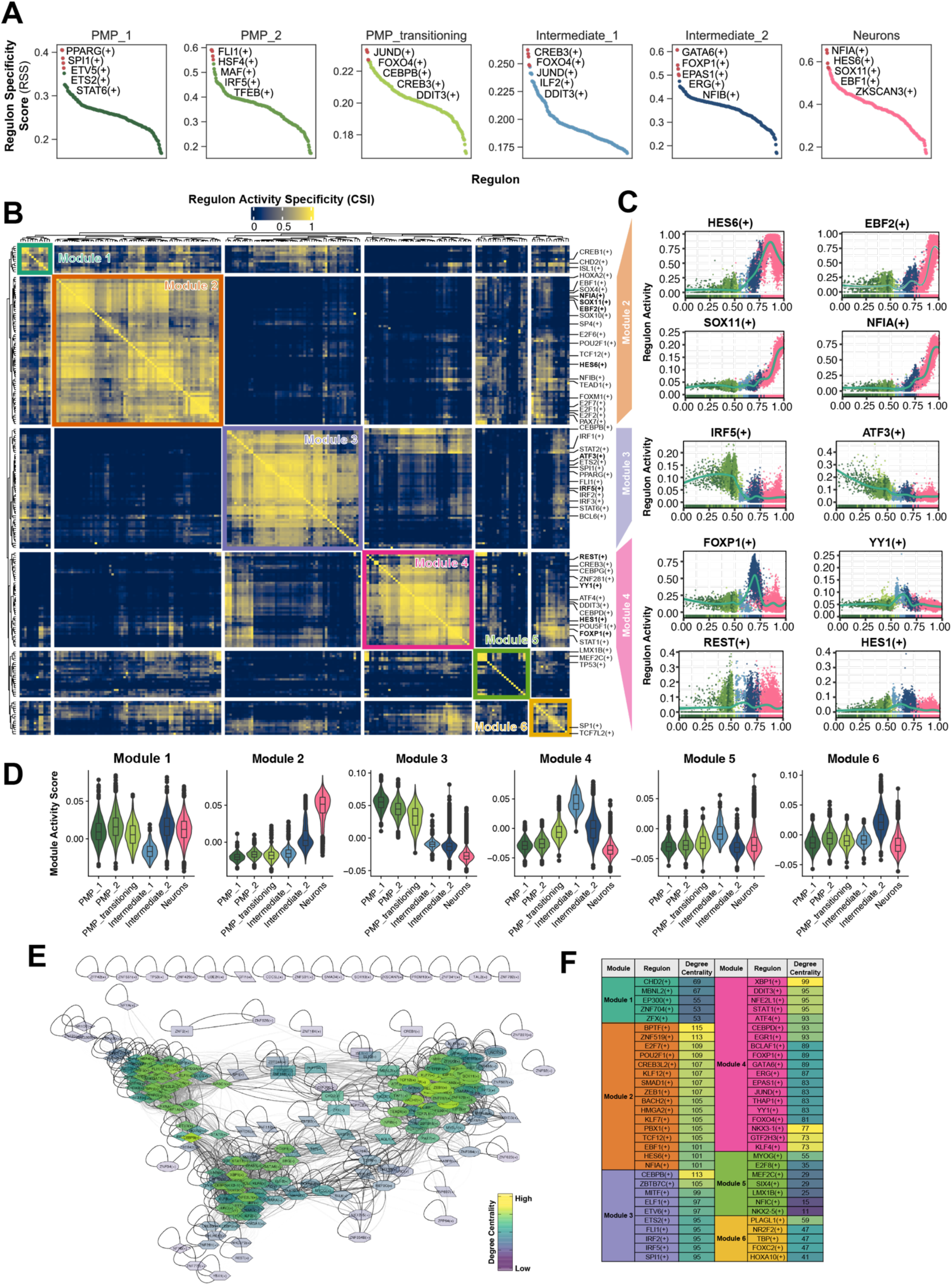
*De novo* regulon and gene regulatory network (GRN) analysis during NEUROG2-mediated reprogramming. A. Ranking of regulons across cell types based on regulon specificity scores (RSS). B. Heatmap showing regulon modules identified by hierarchical clustering of the Connection Specificity Index (CSI) matrix. C. Dynamic plots showing activity changes of selected key regulons along the pseudotime trajectory. D. Violin plots showing regulon module activity scores across annotated cell types. E. Regulon association network constructed using CSI-based interactions. Nodes represent regulons and edges denote high-specificity regulatory associations, hub regulons are colored according to degree centrality. F. Table listing top hub regulons within each module based on hubness (degree centrality) scores.

### Human PMP-derived microglial lineage cells undergo neuronal reprogramming *in vivo* following NEUROG2 induction

To determine whether NEUROG2-mediated neuronal reprogramming of human microglial lineage cells can occur *in vivo*, we transplanted human Dox-inducible NEUROG2 PMPs into mouse brains at postnatal day 0 (P0) to generate human microglial chimeric brain models ^49^. Consistent with prior studies from our group and others, transplanted cells do not give rise to human neurons under baseline conditions, owing to the differentiation protocol that yields a highly pure population of PMPs ^47,48,57,58^. Accordingly, NEUROG2 expression was induced through Dox administration at defined post-engraftment time points to assess *in vivo* MtN conversion (Fig. 4A). At six weeks following Dox administration (corresponding to a mouse age of approximately three months), engrafted human cells expressing human nuclear antigen (hN) were readily detected in the mouse brain, a subset of which expressed NEUROG2 (Fig. 4B, 4E). Transplanted human cells (hN⁺) were readily detected across multiple brain regions, including the corpus callosum (CC) overlying the hippocampus, where DCX⁺ immature neurons are normally absent in the adult brain, subventricular zone (SVZ) and the olfactory bulb (OB), a region that physiologically supports the generation of DCX⁺ immature neurons (Fig. 4C). Importantly, while the majority of engrafted human cells differentiated into hN⁺/IBA1⁺ microglia, a subset exhibited neuronal fate conversion, evidenced by co-expression of the immature neuronal marker DCX together with the human marker (DCX⁺ hN⁺; Fig. 4C). DCX⁺ hN⁺ cells were observed in the OB, SVZ and CC, indicating that the reprogramming response was not confined to a single niche but could occur across anatomically distinct environments (Fig. 4C). High-magnification views and 3D-reconstruction imaging revealed DCX⁺ hN⁺ cells with elongated morphologies and polarized processes consistent with neuronal-like remodeling in situ (Fig. 4C, inset/zoom). In contrast, in animals maintained without Dox administration, transplanted human cells were still detectable, but DCX co-labeling was not observed in either OB or CC at the same 3-month time point (Fig. S3), supporting that neuronal marker induction depended on Dox-triggered NEUROG2 activation rather than nonspecific effects of transplantation or engraftment. Quantification across animals demonstrated that Dox treatment produced a measurable fraction of human graft-derived cells acquiring neuronal identity, with approximately ∼15–20% of hN⁺ cells retaining NEUROG2 signal at the analyzed time point and ∼7–10% of hN⁺ cells expressing DCX (Fig. 4E). These data indicate that NEUROG2 induction can be achieved in a subset of grafted cells *in vivo* and is accompanied by neuronal marker acquisition in a defined fraction of the human microglial lineage population. Together, these findings provide *in vivo* proof-of-concept that xenografted human microglial lineage cells can be induced to adopt neuronal identity in the adult brain environment after NEUROG2 induction.

**Fig 4.**
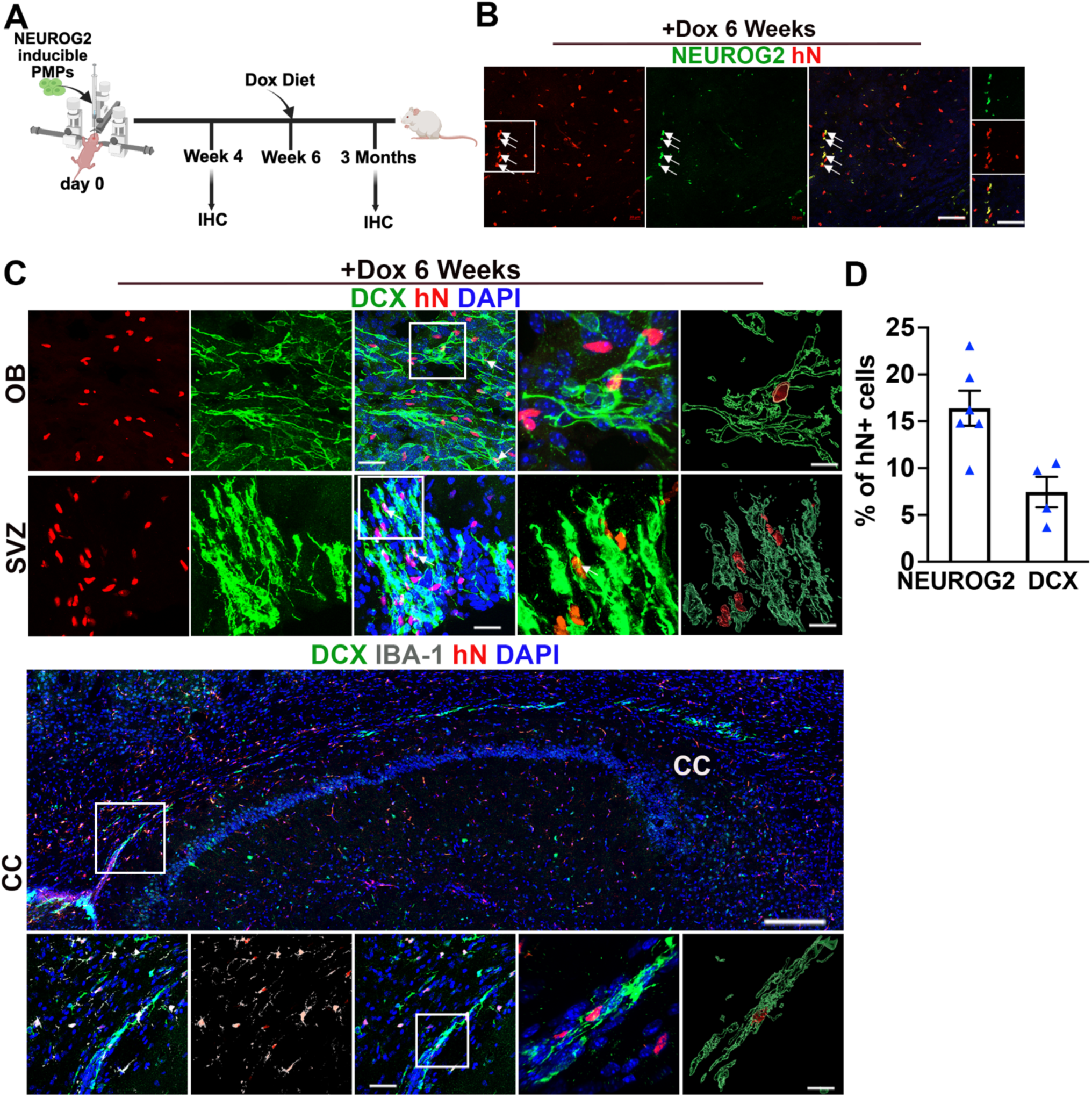
NEUROG2-induced human microglial lineage cells engraft into the mouse brain and acquire neuronal markers following Dox induction. A. Experimental design. B. Representative images of NEUROG2 (iNGN2) and hN transplanted cells. Scale bars, 50 μm and 20 μm in the original and enlarged images, respectively. C. Representative images of DCX, IBA-1, and hN transplanted cells at 6 weeks after Dox induction. Representative raw fluorescence super-resolution and 3D surface rendered images showing colocalization of DCX^+^ hN^+^ staining. Arrows indicate DCX^+^hN^+^ cells. Scale bars, 200, 20, and 5 μm. D. Quantification of neuronal marker acquisition among engrafted human cells at 6 weeks post-induction. Data are presented as mean ± SEM; individual values represent the number of mice.

## Discussion

Direct lineage reprogramming represents a promising approach to generate neurons independently of endogenous neurogenesis or exogenous neuronal transplantation. However, substantial uncertainty remains regarding lineage specificity, reproducibility, and the translatability of direct reprogramming approaches to human systems. In the Waddington epigenetic landscape^59^, microglia occupy a deeply stabilized myeloid basin, long thought to preclude neuronal conversion. Our findings demonstrate that NEUROG2 overcomes this constraint in human microglial lineage cells, enabling direct neuronal conversion in vitro and in chimeric brains.

While macroglia, including astroglia and oligodendroglia, have been shown to be amenable to neuronal reprogramming, a long-standing and unresolved question in the field has been whether myeloid-derived microglial cells can be redirected toward a neuronal fate. Here, we provide convergent lines of evidence demonstrating that human microglial lineage cells, specifically hESC-derived microglial populations, can indeed transition toward a neuronal identity following activation of NEUROG2, despite their profound developmental and epigenetic divergence from neuroectoderm-derived neural cells. First, using a rigorously defined human microglial lineage differentiation platform to generate highly pure human microglial cells, our snRNA-sequencing analyses delineate the molecular logic of this transition. Rather than an abrupt fate switch, microglial reprogramming proceeds along a continuous and ordered transcriptional trajectory comprising sequential stages: (1) suppression of immune lineage programs, (2) engagement of extracellular matrix remodeling and structural reorganization pathways, and (3) emergence of transcriptional networks supporting neuronal signaling, calcium handling, axonogenesis, and synaptic function. This progression is independently supported by pseudotime ordering, RNA velocity, and CytoTRACE analyses, all of which converge on a unidirectional lineage transition. The identification of discrete transcription factor regulatory modules further indicates that conversion is governed by structured regulatory cascades rather than nonspecific global transcriptional activation. Second, electrophysiological and immunocytochemical analyses provide compelling functional and structural validation, showing that NEUROG2-converted human microglial lineage cells acquire hallmark properties of excitable neurons and exhibit synaptic protein organization. Third, human microglia chimeric brain models provide a stringent and definitive system for *in vivo* lineage tracing ^49^. By transplanting highly pure hiPSC-derived PMPs into the mouse brain, any resulting human neurons can be unequivocally attributed to the reprogrammed grafted cells, as the host brain contains no endogenous human neurons. This approach eliminates background ambiguity and establishes an ideal platform for rigorously testing the feasibility and fidelity of in vivo MtN reprogramming from xenografted human microglia. Using this system, we demonstrate that xenografted human microglia can be converted into neurons *in vivo* and that the induced neuronal identity is not transient. Following engraftment and *in vivo* induction of NEUROG2 expression, a subset of human cells acquires neuronal marker expression across different brain regions. Altogether, these findings provide definitive evidence that human microglial lineage cells can be reprogrammed into neurons by NEUROG2.

Our findings substantially extend prior work in this area, as all previous studies of microglia- and macrophage-to-neuron conversion have been confined to rodent systems and have produced inconsistent and often conflicting results ^25,30,31^. Given the pronounced species differences between human and mouse microglia, particularly in transcriptional identity and regulatory networks, MtN strategies and conversion mechanisms defined in rodents may not be directly applicable to human microglia. By demonstrating robust acquisition of neuronal identity in human microglial lineage cells, our study provides direct evidence that human microglia are likewise amenable to neuronal fate conversion. The use of hPSC-derived microglial lineage cells as neuron-generating vehicles is therefore highly translationally relevant. Neurodegenerative diseases and CNS injuries are invariably accompanied by pronounced neuroinflammation, a process largely initiated and orchestrated by microglia. Under physiological conditions, microglia are essential for maintaining brain homeostasis and supporting neuronal integrity. In aging-associated neurodegenerative states, however, microglia undergo maladaptive transitions marked by senescent or dystrophic features, loss of neuroprotective capacity, pronounced morphological alterations, dysregulated lipid metabolism, and pathogenic transcriptional programs that actively drive neurodegeneration and cognitive decline ^60–64^. Our recent work ^48^, together with studies from others ^57,65–67^, has shown that transplantation of healthy, functionally competent human microglia or microglia-like cells can restore neuroprotective functions and remodel the diseased brain environment, underscoring their therapeutic potential. Building on this foundation, MtN reprogramming offers a dual-function strategy in which transplanted microglia not only modulate a hostile neural microenvironment but also serve as carriers of inducible reprogramming factors, enabling their conversion into neurons in situ. In addition, microglia possess a unique capacity for extensive migration following transplantation ^35–38^. By exploiting this intrinsic dispersal property, microglia engineered with inducible reprogramming programs could populate broad regions of the brain, and upon controlled activation, generate neurons in a spatially distributed and disease-relevant manner.

While the findings reported here are compelling, further studies are warranted to assess the functional maturation and synaptic integration of the converted neurons in vivo within chimeric brain environments. More critically, it will be essential to evaluate human MtN conversion and neuronal function in disease-relevant and human-specific pathological contexts. Although we achieved robust reprogramming efficiency *in vitro* and established proof-of-concept evidence for human MtN conversion *in vivo*, the overall conversion efficiency in vivo remains relatively low. This limitation may be addressed through strategies such as AAV-mediated enhancement of NEUROG2 expression ^68^ or combinatorial approaches incorporating small-molecule modulators ^69^ to further optimize reprogramming efficiency and neuronal maturation. In summary, this study demonstrates that human microglial lineage cells can be redirected toward a neuronal identity and systematically defines the transcriptional, structural, and functional features of this process. By integrating human cellular platforms, transcriptomic trajectory analyses, functional validation, and in vivo assessment, our findings advance understanding of human MtN plasticity and provide a strong foundation for future mechanistic and translational investigations.

## Acknowledgements

This work was in part supported by grants from the NIH (R01NS122108 and R01AG073779 to P.J.). M.J. was supported by a postdoctoral fellowship award from the New Jersey Department of Health (CAUT24DFP004). Y.L. was supported by NIH R01NS110707. NIH (R01AG064579 and RF1NS128800), the JSRM Foundation, BrightFocus Foundation (BFF17-0008), and the Alzheimer’s Association to S.F. We thank Dr. James Knowles’s group at Rutgers University and the Human Genetics Institute of New Jersey for providing facilities and assistance with snRNA-seq library preparation and live-cell imaging.

## Author Contributions

M.J. and P.J. designed the experiments and interpreted the data; M.J. carried out most of the experiments with technical assistance from H.Z. and R.D.; Z.M. performed RNA-seq data analyses and assisted with the interpretation of sequencing data. R.D. performed electrophysiological recordings; Y.L. and H.X. designed and performed gene editing in the hESC lines; S.F. provided critical suggestions to the study and assisted with imaging data analysis. P.J. conceived the study, directed the project, and co-wrote the manuscript with M.J. and Z.M., with input from all co-authors.

## Competing Interests

The authors declare no competing interests.

## Methods

### Targeting vector pUCM-AAVS1-TO-hNGN2

The pUCM-AAVS1-TO-hNGN2 plasmid was obtained from Addgene (cat. no. 105840). The targeting vector contains the following elements (Fig. S1A): a 5′ homology arm (AAVS1 5′-HA), a splice acceptor (SA), a T2A self-cleaving sequence, a puromycin resistance cassette, a bovine growth hormone (BGH) poly(A) signal, an EF1α promoter driving the mCherry cassette followed by a BGH poly(A) signal, a CAG promoter driving the rtTA cassette, a TRE3G element, the hNGN2 cDNA, an SV40 poly(A) signal, and a 3′ homology arm (AAVS1 3′-HA). To minimize potential counter-regulatory or promoter-interference effects, the hNGN2 cDNA cassette was positioned in the opposite orientation relative to the TRE3G promoter. All fragments were assembled by Gibson assembly (New England Biolabs) as described previously (Gibson et al., 2009; Li et al., 2017) The final constructs were selected with ampicillin. Following TALEN-mediated gene editing with pZT-AAVS1-L1 (Addgene cat. no. 52637) and pZT-AAVS1-R1 (Addgene cat. no. 52638), positively targeted cells were identified by constitutive mCherry expression. Upon doxycycline treatment, hNGN2 expression was induced within 48 hours, thereby driving neuronal reprogramming.

### Construction of pUCM-CLYBL-TO-hNGN2 targeting vector

To generate the pUCM-CLYBL-TO-hNGN2 targeting vector, pUCM-AAVS1-TO-hNGN2 (Addgene, cat. no. 105840) was modified as follows: (1) The EF1α promoter–driven mCherry cassette was removed. (2) An IRES-EGFP cassette was inserted immediately downstream of rtTA, enabling constitutive EGFP expression under the CAG promoter. (3) Core insulator sequences (232 bp) were excised from HR130PA-1 (System Biosciences) and cloned to flank the targeting cassette. (4) The modified insert generated in steps 1–3 was subsequently cloned into the backbone of pUCM-CLYBL-hNIL (Addgene, cat. no. 105841). The final pUCM-CLYBL-TO-hNGN2 targeting vector therefore consists of the following elements (Fig. S1B): a 5′ homology arm (CLYBL 5′-HA), a splice acceptor (SA), a puromycin resistance cassette, a bovine growth hormone (BGH) poly(A) signal, a core insulator, a CAG promoter driving rtTA, an IRES-EGFP cassette, a β-globin poly(A) signal, a TRE3G element, the hNGN2 cDNA, an SV40 poly(A) signal, a second core insulator, and a 3′ homology arm (CLYBL 3′-HA). To minimize potential counter-regulatory or promoter-interference effects, the hNGN2 cDNA cassette was positioned in the opposite orientation relative to the TRE3G promoter. Following TALEN-mediated gene editing with pZT-C13-L1 (Addgene cat. no. 62196) and pZT-C13-R1 (Addgene cat. no. 62197), positively targeted cells were identified by constitutive EGFP expression. Upon doxycycline treatment, hNGN2 expression was induced within 48 hours, thereby driving neuronal reprogramming.

### Generation of human PSCs

All human PSC lines were maintained in chemically defined mTeSR plus medium (Stemcell Technologies Inc.) in a feeder free fashion and passaged every 4∼5 days at a 1:4∼1:6 ratio using ReLeSR (Stemcell technologies) as described previously ^70,71^. Alternatively, cells were passaged every 4∼5 days with Accutase (Innovative cell technologies) onto Matrigel (BD Biosciences)-coated culture plates at a ratio of 1:4∼1:8 supplemented with ROCK inhibitor Y-27632 (10 mM, R&D systems). To monitor and ensure the genetic stability of the cells, routine karyotyping examination was done every 10 to 15 passages. H1 and H9 hESC lines were used as parental lines to generate a doxycycline-inducible NGN2 H1 hESC line and an inducible NGN2 H9 line carrying the CSF2RB A455D mutation ^48,72^. Approximately 1×10^5^ hESCs were plated on a well of a 12-well plate in mTeSR plus medium, one day prior to transfection. Cells from each well were co-transfected with the vectors (1 µg) and both TALENs plasmids for AAVS1 or CLYBL (left and right TALENs, 0.5 µg each) vectors using Lipofectamine Stem (Life Technologies) following the manufacturer’s instructions. ROCK inhibitor (10 μM) was added for the first 24 h post-transfection to enhance single cell survival. Cells were fed with fresh mTeSR plus medium every day until Day 5 post-transfection, when Puromycin (0.75-1 µg/mL) selection was started. Resistant single clones were isolated and manually picked after two to four days of selection and subsequently expanded individually. Genomic DNA extracted from single clones was examined by PCR for the identification of correctly targeted hPSC clones. Primer sequences for detecting positive clones are listed in Table S1 and Table S2. The PCR products with the correct size were verified by Sanger sequencing.

### Differentiation and culture of PMPs

PMPs were generated from the hESC cell lines following previously established protocols ^50^. Yolk sac embryoid bodies (YS-EBs) were first induced by culturing hESCs in mTeSR 1 media (STEMCELL Technologies) supplemented with bone morphogenetic protein 4 (BMP4, 50 ng/ml), vascular endothelial growth factor (VEGF, 50 ng/ml), and stem cell factor (SCF, 20 ng/ml) for 7 days. To promote myeloid lineage commitment, the YS-EBs were subsequently transferred to X-VIVO 15 medium (Lonza) supplemented with interleukin-3 (IL-3, 25 ng/ml) and macrophage colony-stimulating factor (M-CSF, 100 ng/ml) ^73^. At 4-6 weeks after plating, human PMPs were continuously released into the supernatant and could be harvested for over 3 months.

### Animals and cell transplantation

All animal procedures were conducted without sex bias and were approved by the Rutgers University Institutional Animal Care and Use Committee. PMPs were collected from culture supernatant and resuspended in PBS at 100,000 cells/µl and stereotactically transplanted into P0 Rag2^−/−^ hCSF1 knock-in immunodeficient mice (C;129S4-*Rag2^tm^*^1^*^.1Flv^ Csf1tm1*^(CSF1)^*^Flv^ Il2rg^tm^*^1^*^.1Flv^/J*, The Jackson Laboratory). Bilateral injections were performed at ±1.0 mm lateral to the midline, −2.0 mm posterior to bregma, and at dorsoventral depths of −1.5 and −1.2 mm. Neonatal mice were briefly cryoanesthetized on ice for 4–5 minutes, and 0.5 µl of cell suspension was delivered per site (four sites total) using a digital stereotaxic apparatus (David Kopf Instruments) equipped with a neonatal adapter (Stoelting). Mice were weaned at three weeks of age and maintained for subsequent analyses at designated time points.

### RNA isolation and quantitative reverse transcription PCR

Total RNA was isolated using TRIzol reagent (Thermo Fisher Scientific, 15596026), and complementary DNA (cDNA) was synthesized from 600 ng RNA using the TaqMan™ Reverse Transcription Reagents (Thermo Fisher Scientific; N8080234). Real-time quantitative PCR was performed on an ABI 7500 Real-Time PCR System using TaqMan Fast Advanced Master Mix (Thermo Fisher Scientific). All primers are listed in Table S3. Relative gene expression was calculated using the 2^−ΔΔ*Ct*^ method after normalization to GAPDH.

### Time-lapse imaging

To evaluate morphological changes, PMP cells were seeded on growth factor-reduced Matrigel (Corning) coated in a six-well plate at a density of 2 × 10^5^ cells and allowed to adhere for 12 h prior to Dox treatments using neuron differentiation medium, which is composed of a 1:1 mixture of Neurobasal (Thermo Fisher Scientific) and DMEM/F12 (Hyclone), supplemented with 1x N2, 1x B27-RA (Thermo Fisher Scientific), BDNF (20 ng/ml, Peprotech), GDNF (20 ng/ml, Peprotech), dibutyryl-cyclic AMP (1mM, Sigma), and L-Ascorbic Acid (200 nM). Immediately following treatment, live cell images across 25 regions in one of the six wells were automatically captured every 4 hours for 14 days using a BioTek Cytation C10 Confocal Imaging Reader (BioTek) equipped with temperature- and CO_2_-controlled environmental chambers. Images taken by Cytation C10 were at 10x magnification.

### Electrophysiology

The detailed procedures and analysis for cultured neurons were described previously ^74^. The recording external solution contained (in mM): 140 NaCl, 5 KCl, 10 HEPES, 2 CaCl_2_, 2 MgCl_2_, 10 Glucose, pH 7.4. All cell culture recordings were conducted at room temperature. For whole-cell recordings, the neurons were recorded with glass pipettes (5-8 MΩ) filled with the intracellular solution containing (in mM) 126 K-gluconate, 4 KCl, 10 HEPES, 0.05 EGTA, 4 ATP-magnesium, 0.3 GTP-sodium, and 10 phosphocreatine. The current injection recording was performed with injected current steps (from -50 to 130 pA, 10 pA steps). The voltage-dependent sodium and potassium current recordings were performed under consecutive depolarizing voltage steps from -10 mV to +40 mV (10 mV per step, lasting 250 ms). All data acquisition and analysis were done using pCLAMP 10.7 (Molecular Devices).

### Tissue and coverslip immunostaining, image acquisition, and analysis

Mouse brains and coverslips were fixed with 4% paraformaldehyde. The mouse brains were placed in 20% and later in 30% sucrose for dehydration. Following dehydration, brain tissues were immersed in OCT and frozen for sectioning. The frozen tissues were cryo-sectioned with 30-μm thickness for immunofluorescence staining. The tissues were blocked with a blocking solution (5% goat serum in PBS with 0.8% Triton X-100) at room temperature (RT) for 1 hour. The coverslips were blocked with a blocking solution (5% Goat Serum + 1% BSA). The primary antibodies were diluted in the same blocking solution and incubated with the tissues or coverslips at 4 °C overnight (all the primary antibodies are listed in Table S4). The sections were washed with PBS and incubated with secondary antibodies for 1 hour at RT. After washing with PBS, the slides were mounted with anti-fade Fluoromount-G medium containing 1, 40,6-diamidino-2-phenylindole dihydrochloride (DAPI) (Southern Biotechnology). All images were captured with a Zeiss 800 confocal microscope.

### Single-nucleus preparation and snRNA-seq library construction

Single nuclei were prepared using Nuclei PURE Prep kit (Sigma) following the manufacturer’s protocols. Isolated nuclei suspensions were loaded onto the Chromium Controller for snRNA-seq library preparation. Libraries from 12-day samples were generated using the Chromium Next GEM Single Cell 3′ Kit v3.1, 5-day samples were processed using the Chromium GEM-X Single Cell 3′ Kit v4 in combination with the Dual Index Kit TT Set A, following the manufacturer’s protocols. Library quality and fragment size distributions were assessed using an Agilent 2100 Bioanalyzer. Sequencing was performed on an Illumina NovaSeq 6000 system using S4 flow cells.

### Single-nucleus RNA sequencing data analysis

Raw sequencing reads were aligned to the human GRCh38 reference genome using Cell Ranger (v9.0.1). Genes expressed in fewer than 20 cells were excluded from downstream analyses. Nuclei were retained if they contained between 500 and 20,000 detected genes and exhibited mitochondrial transcript content below 5%. Potential doublets and multiplets were identified using Scrublet ^75^ but were not excluded from subsequent analyses. Downstream analyses were conducted using Scanpy ^76^ (v1.11.5) with Python v3.10.19. After normalization and log transformation, 2,000 highly variable genes (HVGs) were identified and used for principal component analysis (PCA). Batch effects were corrected using Harmony ^77^, which converged after two iterations. Nonlinear dimensionality reduction was performed using Uniform Manifold Approximation and Projection (UMAP) based on the top 50 principal components, with 50 nearest neighbors and a minimum distance parameter of 0.2. Graph-based clustering was performed using the Leiden algorithm with a resolution of 0.5, resulting in the identification of seven clusters.

Volcano plot was produced using EnhancedVolcano R package (v1.28.2) with R v4.5.2. Differentiation potential was inferred using the CytoTRACE ^78^ kernel imported from CellRank 2 ^79^ python package (v2.0.7), CytoTRACE scores were computed from normalized expression matrices and used to assess relative differentiation potential across cell populations. For RNA velocity analysis, spliced and unspliced transcript count matrices were generated using the velocyto.py ^80^ package (v0.17.17). RNA velocity and velocity pseudotime were subsequently inferred using scVelo ^81^ (v0.3.3). Dynamic gene expression changes along pseudotime were visualized using heatmaps generated with the SCP/scop R package. Gene ontology (GO) enrichment analysis was performed using the clusterProfiler ^82^ R package (v4.18.2).

Gene regulatory networks were inferred using the pySCENIC ^83,84^ framework (v0.12.1). Regulon modules were defined based on the Connection Specificity Index (CSI), a context-dependent metric that quantifies the specificity of associations between regulons ^83,85^. Briefly, Pearson correlation coefficients (PCCs) were calculated for all pairs of regulon activity scores. For a given pair of regulons, A and B, the CSI was defined as the fraction of regulons whose PCC with both A and B was lower than the PCC observed between A and B. Hierarchical clustering using Euclidean distance was applied to the CSI matrix to identify distinct regulon modules, which were visualized using the ComplexHeatmap ^86,87^ R package. Regulon activity scores were calculated using AddModuleScore() function from Seurat ^88^ R package (v5.3.1). To further characterize regulon interactions, a regulon association network was constructed using a CSI threshold of >0.7. Network visualization and analysis were performed in Cytoscape ^89^ (v3.10.1). Hub regulators were identified based on degree centrality, calculated using the cytoHubba ^90^ (v0.1) plugin.

### Statistical analysis

All data are represented as mean ± SEM. When only two independent groups were compared, significance was determined by using a two-tailed unpaired t-test with Welch’s correction. A p-value of < 0.05 was considered significant. All analyses were performed in GraphPad Prism v.9. All experiments were independently performed at least 3 times with similar results.

## Data availability

The GEO accession number of the single-nuclear RNA sequencing data reported in this study is GSE316615, with reviewer access token *mtkrucimhvejdev*.

**Video S1 and S2:** Longitudinal time-lapse imaging of the reprogramming process demonstrates that, over several days, NEUROG2-induced cells progressively extend neurite-like processes (Video S1), a dynamic morphological change that is absent in cultures without doxycycline treatment (Video S2).

## Figure legends

**Figure S1.**
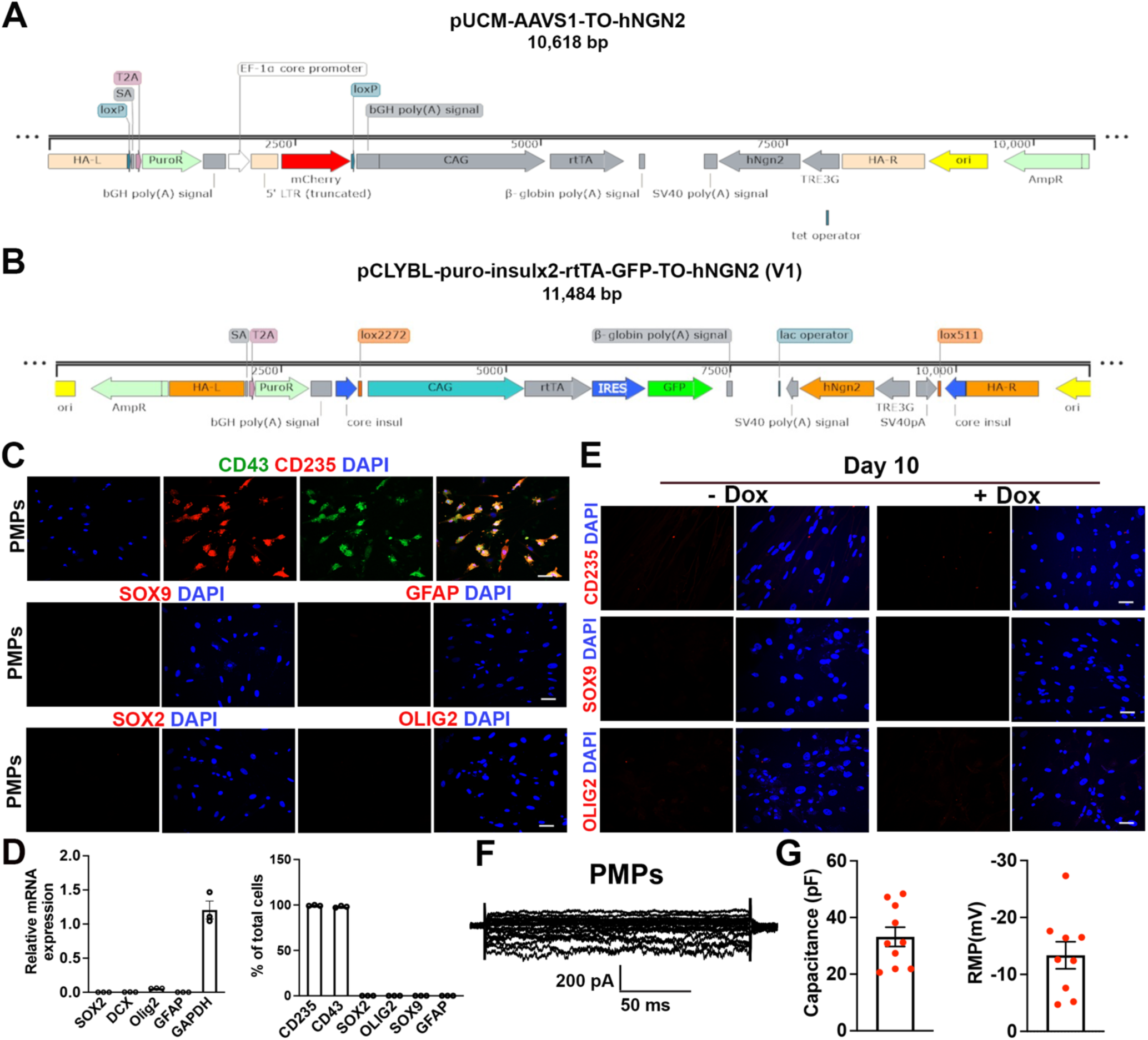
Human iPSC-derived primitive macrophage progenitors exhibit a pure myeloid identity and lack neural lineage markers prior to reprogramming. A. Schematic of the AAVS1-targeting donor construct pUCM-AAVS1-TO-hNGN2 used to generate inducible NEUROG2 human embryonic stem cell (hESC) lines. The construct contains a tetracycline-responsive element (TRE3G) driving human NEUROG2, a CAG-driven reverse tetracycline transactivator (rtTA), and selection cassettes flanked by homology arms (HA-L and HA-R) for targeted integration into the AAVS1 locus. B. Schematic of the lentiviral construct pCLYBL-puro-insulx2-rtTA-GFP-TO-hNGN2 (V1) used for inducible NEUROG2 expression. The construct includes a CAG-driven rtTA-IRES-GFP cassette, a TRE3G-regulated NEUROG2 expression cassette, and selection markers enabling doxycycline-dependent transgene induction. C. Representative images of CD43, CD235, SOX9, GFAP, SOX2, and OLIG2. Scale bars, 20 μm. D. Quantification of lineage marker expression. Data represent mean ± SEM from biological replicates. E. qRT-PCR analysis of neural lineage transcripts (SOX2, DCX, OLIG2, GFAP) in PMP cultures. Expression levels are normalized to GAPDH and presented as mean ± SEM. F. Representative images of SOX9, OLIG2, and CD235. Scale bars, 20 μm. G. Representative voltage-clamp recordings from untreated PMP cells showing small, non-neuronal membrane currents, consistent with a non-excitable myeloid phenotype. H. Quantification of passive membrane properties of PMP cells, including membrane capacitance and RMP. Data are presented as mean ± SEM; each point represents an individual recorded cell.

**Figure S2.**
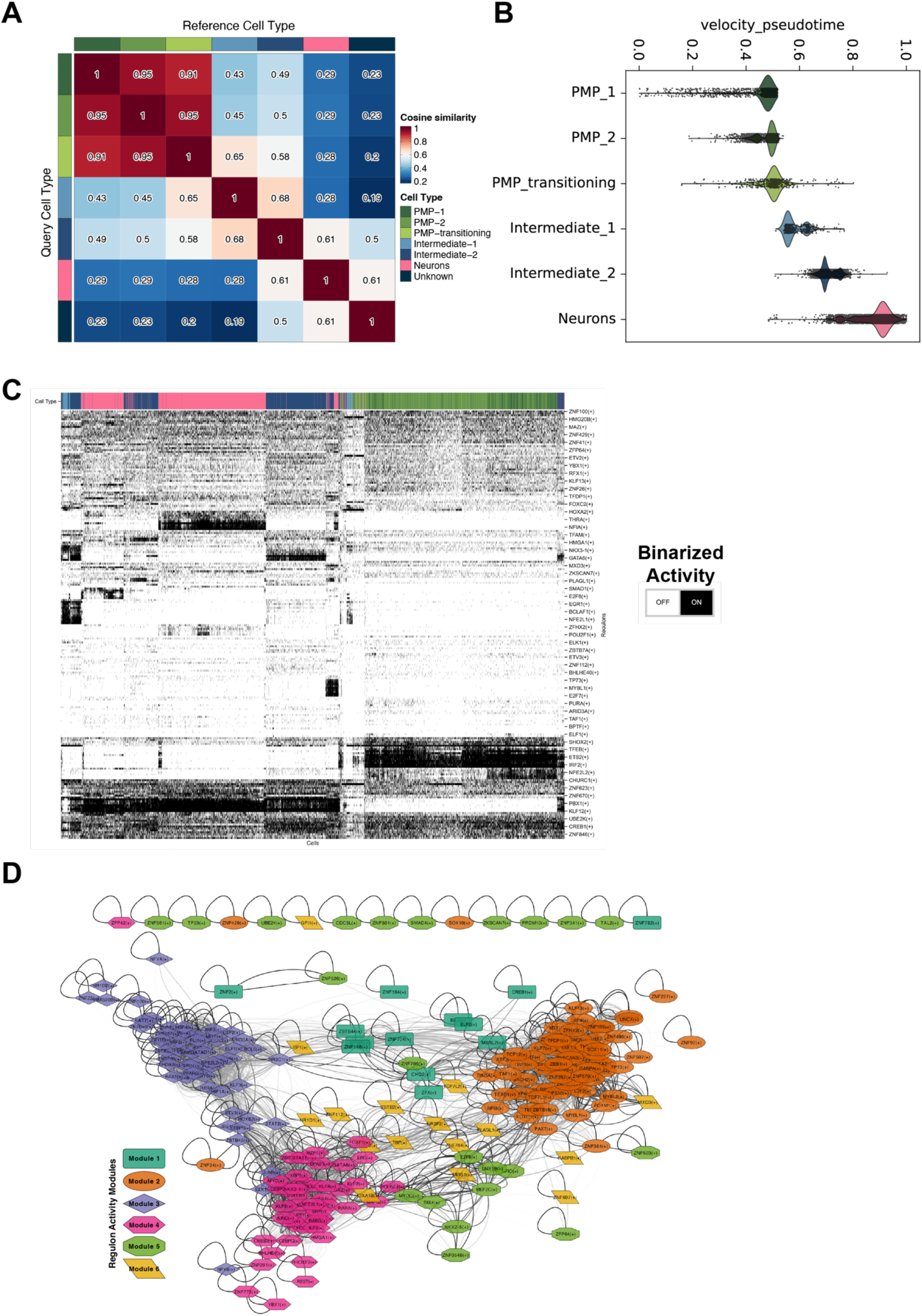
NEUROG2 induction drives coordinated transcriptional state transitions and reorganization of gene regulatory networks during reprogramming. A. Heatmap showing cosine similarity indices comparing 7 identified Leiden clusters (cell types). B. Violin plot showing velocity pseudotime distribution across major cell types. C. Heatmap showing binarized regulon activity states (active/inactive) inferred using the SCENIC pipeline. D. Regulon association network constructed using CSI-based interactions colored by modules.

**Figure S3.**
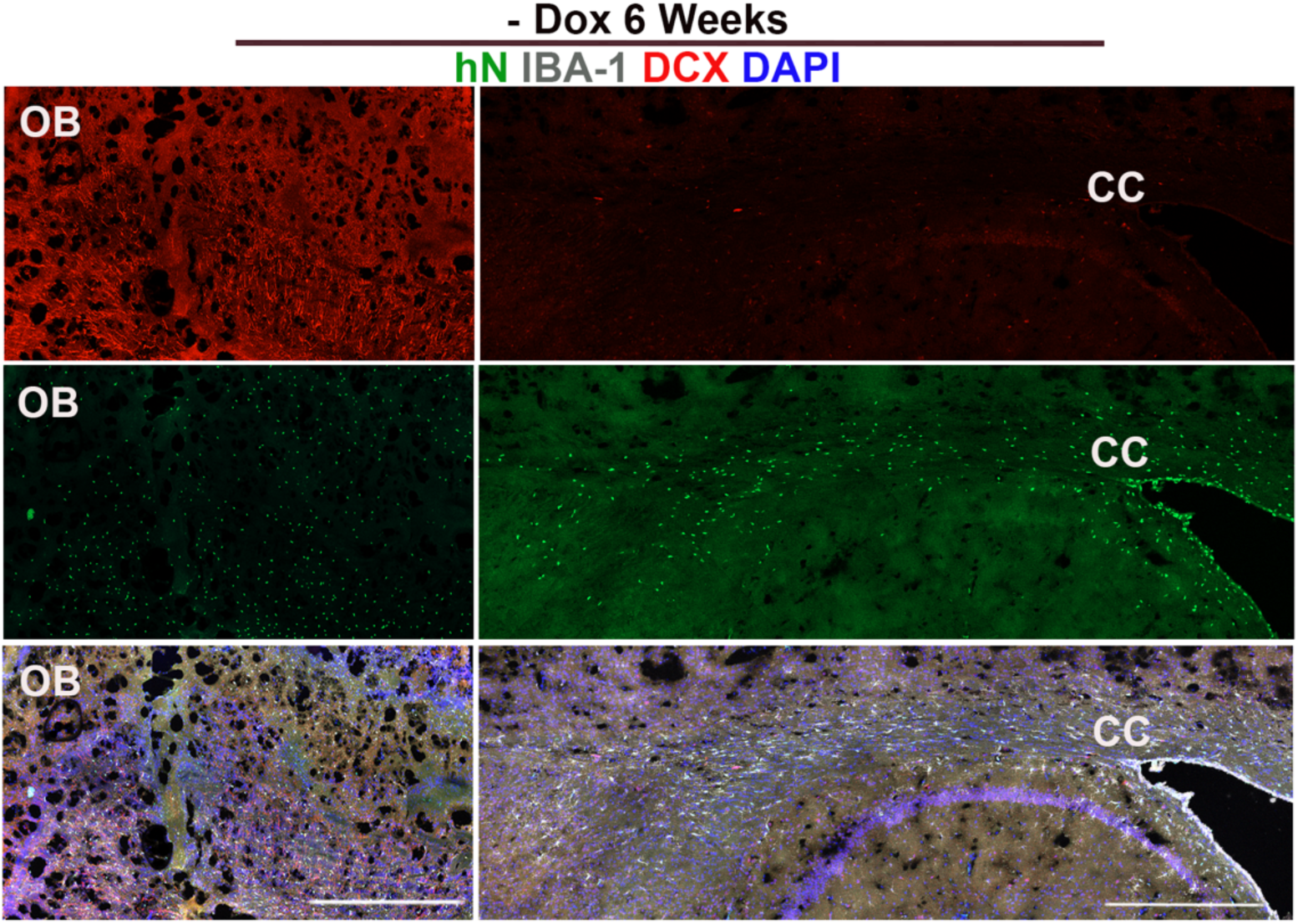
Absence of neuronal marker induction in xenografted human microglial lineage cells without Dox treatment. Representative images of DCX, IBA-1, and hN transplanted cells at 3 months without Dox induction. Scale bars, 500 μm.

## Supplementary Tables

**Table S1.**
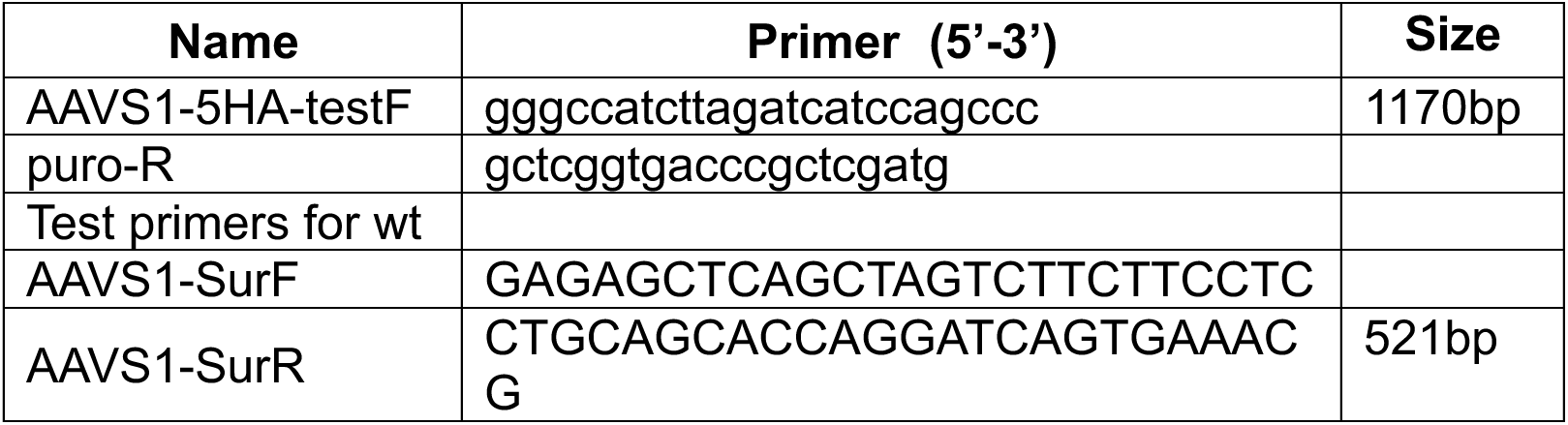
Test primers for gene editing of pUCM-AAVS1-TO-hngn2(C550).

**Table S2.**
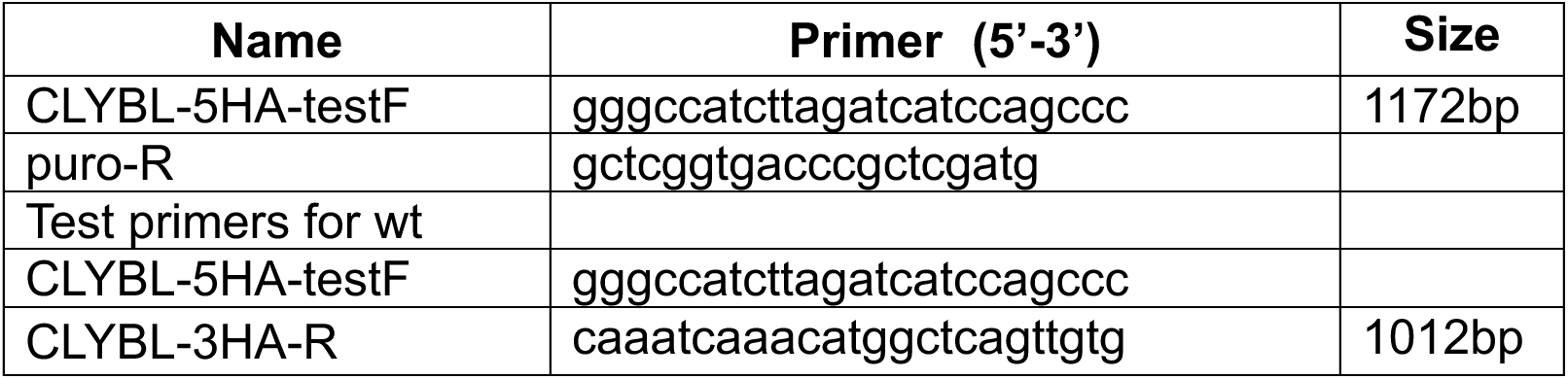
Test primers for gene editing of pUCM-CLYBL-TO-hngn2(F153).

**Table S3.** Antibodies used in this study.

| Reagent | Source | Catalog no. |
| --- | --- | --- |
| Anti-CD235 | Invitrogen | PA5-27154 |
| Anti-CD43 | Invitrogen | 14-0439-82 |
| Anti-DCX | Cell Signaling Technology | 4604S |
| Anti-GFAP | Millipore Sigma | AB5804 |
| Anti-human Nuclei | Millipore Sigma | MAB4383 |
| Anti-human-TMEM119 | Invitrogen | PA5-62505 |
| Anti-MAP2 | Santa Cruz | sc-74420 |
| Anti-NeuN | Millipore Sigma | ABN78 |
| Anti-Olig2 | Phosphosolutions | 1538 |
| Anti-SOX2 | Millipore Sigma | AB5603 |
| Anti-SOX9 | Cell Signaling Technology | 82630S |
| Anti-TUJ1 | Millipore Sigma | MAB4383 |
| Anti-NEUROG2 | Cell Signaling Technology | 13144S |
| Anti-MAP2 | Invitrogen | PA5-62505 |
| Goat anti-mouse IgG,<br>Alexa Fluor 594 | Invitrogen | A11032 |
| Goat anti-mouse IgG,<br>Alexa Fluor 488 | Invitrogen | A11029 |
| Goat anti-rabbit IgG, Alexa<br>Fluor 488 | Invitrogen | A27034 |
| Goat anti-rabbit IgG, Alexa<br>Fluor 594 | Invitrogen | A11037 |

**Table S4.** Taqman Primers.

| Gene | Gene expression assay catalog number |
| --- | --- |
| *OLIG2* | Hs00300164_s1 |
| *SOX2* | Hs01053049_s1 |
| *DCX* | Hs00167057_m1 |
| *GFAP* | Hs00909233_m1 |
| *GAPDH* | Hs04420697_g1 |

## Notes

### Competing Interest Statement

P.J., M.J., and Y.L. have filed a provisional patent related to this work (Application No.: 63/981,620).

